# A positive feedback loop between germ cells and gonads induces and maintains cnidarian sexual reproduction

**DOI:** 10.1101/2024.05.16.594501

**Authors:** Camille Curantz, Ciara Doody, Helen R Horkan, Gabriel Krasovec, Paris K Weavers, Timothy Q DuBuc, Uri Frank

## Abstract

The fertile gonad includes cells of two distinct developmental origins: the somatic mesoderm and the germline. How somatic and germ cells interact to develop and maintain fertility is not well understood. Here, using grafting experiments and transgenic reporter animals, we find that a specific part of the gonad–the germinal zone–acts as a sexual organizer to induce and maintain *de novo* germ cells and somatic gonads in non-sexual tissue in the cnidarian *Hydractinia symbiolongicarpus*. We further show that germ cells express a novel member of the TGF-β family (*Gonadless*, *Gls*) that induces somatic gonad morphogenesis. Loss of *Gls* resulted in animals lacking somatic gonadal structures but having few, non-proliferative germ cells. We propose a model according to which a small number of primary germ cells drive gonad development though Gls morphogen secretion. The germinal zone in the newly formed gonad, in turn, provides positive feedback to induce secondary germ cells by activating *Tfap2* (the master regulator of germ cell fate) in resident pluripotent stem cells. *De novo* induction of germ cells by gonads in adult life is absent in bilaterians with a sequestered germline. However, the contribution of germ cell signaling to the patterning of somatic gonadal tissue, as observed in *Hydractinia*, may be a general animal feature.

## Introduction

Jointly, germ cells and somatic gonadal cells constitute the main reproductive organ. Their respective developmental origin, however, is spatially and temporally distinct. In most animals, germ cells are generated only once in a lifetime during early embryonic development^1,2^. Somatic gonad tissues develop later and are derived from mesoderm. Once specified, primordial germ cells migrate to the region of developing gonads where they are incorporated with somatic gonad cells^2,3^. Defects in integration result in impaired fertility^4^. Somatic gonad cells communicate with germ cells using multiple signaling pathways to regulate gamete formation^3^. However, the nature of the signals from germ cells to the gonad remains poorly understood. Numerous studies propose that germ cells are dispensable for early embryonic gonad development in mammals but that they are essential later for its proper maturation^5^. In zebrafish, gonads fail to regenerate in the absence of germ cells^6^. In *Drosophila*, germ cells control the differentiation, proliferation and morphogenesis of the somatic follicle cells that surround them by activating the delta/notch signaling pathway^7^. However, data are often contradictory between male and female, or across different species of vertebrates and invertebrates. Furthermore, these studies rely on the ablation of germ cells before their integration in the gonad; this is difficult to perform in animals with maternal germ cell induction. Therefore, the potential implication of an early signaling of germ cells to the gonad somatic tissue to regulate their morphogenesis remains a possibility that requires further studies using other animals^5^. Overall, these processes, being restricted to early embryogenesis, are difficult to access and manipulate in most animal models. To address this issue, we have studied germ cell and soma interactions in the developing and mature gonads of the cnidarian *Hydractinia symbiolongicarpus*, an animal that continuously develops new gonads and induces new germ cells from pluripotent stem cells^8^.

### *Hydractinia* sexual development is continuous

*Hydractinia symbiolongicarpus* is a clonal and colonial cnidarian, a relative of jellyfishes and corals^9^. A *Hydractinia* colony is either male or female and is structured as a network of substratum-attached gastrovascular tubes (called stolons) from which zooids (called polyps) bud continuously as the stolons elongate. The colony contains two primary types of polyps. The first polyp to emerge during metamorphosis of the sexually derived larva is of the feeding type. These polyps possess a head with a mouth surrounded by muscular and innervated tentacles, used to catch prey (Figure 1A). About two months later, when the colony reaches some 100 feeding polyps, a new type of polyp emerges–the sexual polyp. Sexual polyps are the animal’s gonads. The body column of sexual polyps lacks a functional mouth and has only rudimentary tentacles around its oral end. They obtain food through the shared stolonal network. The region below the sexual polyp head (the ‘neck’) is called the germinal zone (Figure1B). Here, adult pluripotent stem cells (known as i-cells^8,10^) are induced to germ cell fate by activation of *Tfap2*, the master regulator of germ cell commitment (Figure1C)^11^. They then migrate to the gastrodermis, proliferate, and move into containers, known as sporosacs, where they mature to gametes (Figure1B). Stolonal elongation, new polyp budding (both feeding and sexual), and germ cell induction from pluripotent i-cells occurs continuously following sexual maturation, making all stages of sexual development available at any one time.

**Figure 1.**
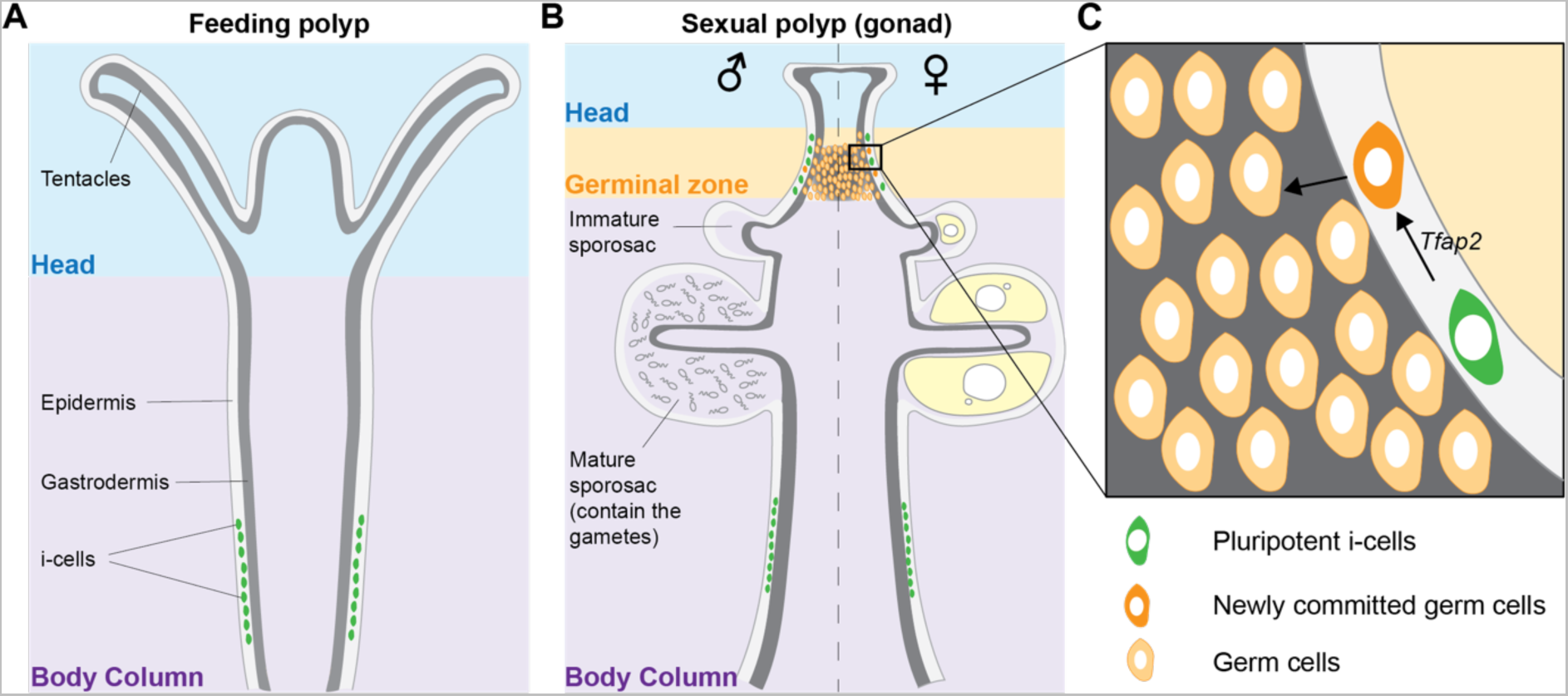
The *Hydractinia* sexual development. (**A**) Feeding polyp morphology showing i-cell localization (green) in the epidermis of the lower part of the body column. (**B**) Sexual polyp morphology showing germ cells (orange) and their commitment from i-cells in the germinal zone. Once committed, germ cells become gastrodermal and start their maturation into functional gametes. The left side represents a male and the right side a female. (**C**) Close up of the germinal zone (black rectangle in B) showing i-cells being induced to germ cell fate by Tfap2 expression.

### The germinal zone is a sexual organizer

To study the maintenance of sexual tissue in *Hydractinia*, we performed regeneration experiments of feeding and sexual polyps in a *Tfap2::GFP* transgenic reporter animal. *Tfap2* is an early germ cell marker; it drives the commitment of pluripotent i-cells to germ cell fate. Feeding polyps of *Tfap2::GFP* reporter animals have no GFP fluorescent cells, while in their sexual polyps, all early germ cells are GFP^+^. Fluorescence dissipates as early germ cells mature to gametes, reflecting *Tfap2* expression in early but not late germ cells^11^ (Figure 2A).

**Figure 2.**
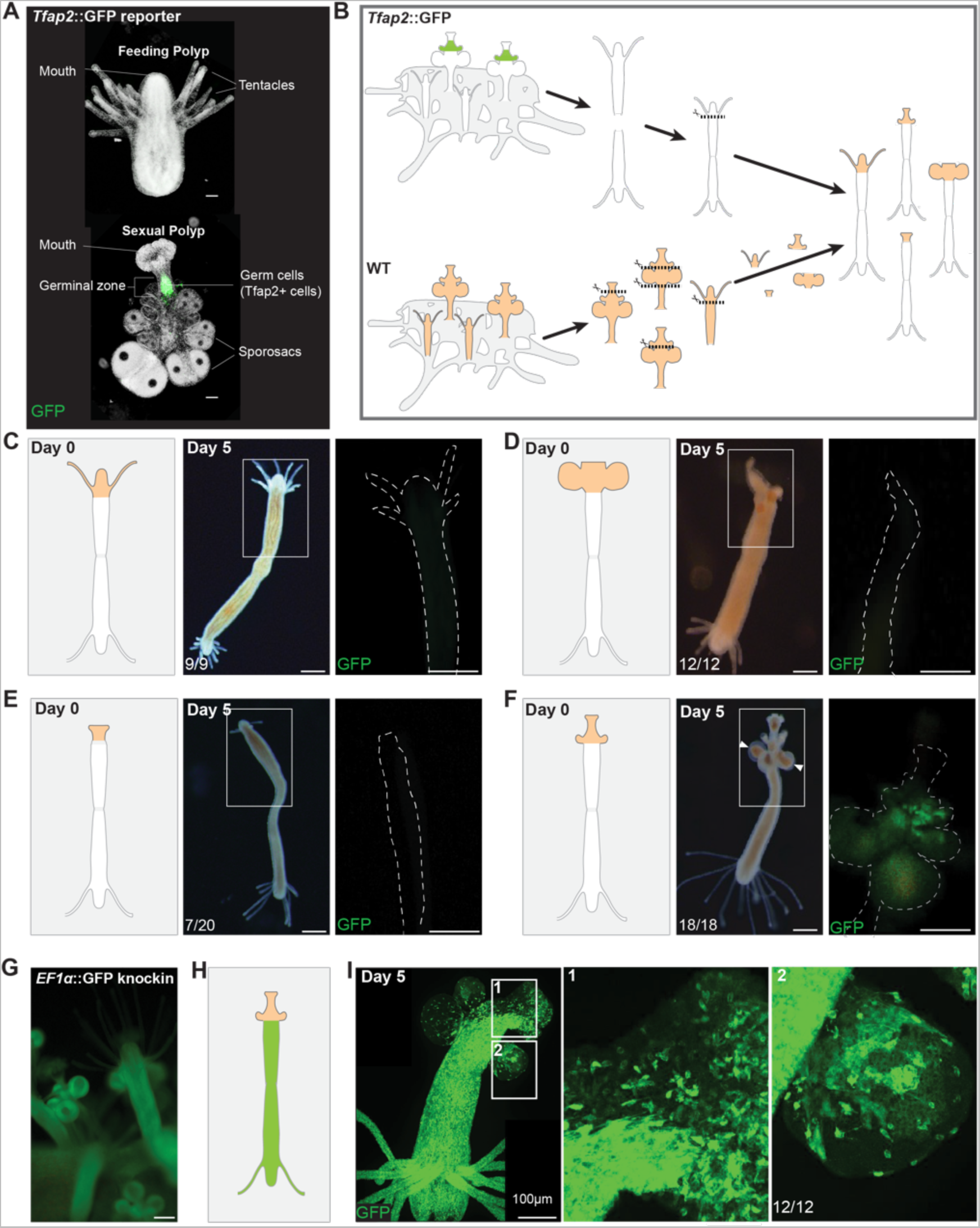
*De novo* induction of germ cells and somatic gonad tissue upon heterotopic grafting of gonad tissue onto feeding polyps. (**A**) *Tfap2*::GFP reporter animal. (**B**) Cartoon of the experimental grafting procedure showing different parts of sexual polyps grafted to a feeding polyp body column. (**C-F**) Outcomes 5 days post grafting of feeding head grafting (C); sporosac region without germinal zone grafting (D); oral tip only (E); head including germinal zone graft showing induction of new germ cells (GFP^+^) in the recipient feeding polyp (F). White arrows show developing sporosacs in F. White dashed line shows the outline of the polyp. (**G**) An *Ef1a*::GFP knock-in animal with ubiquitous GFP expression. (**H**) Cartoon of a chimera between a wild type germinal zone (orange) and an *Ef1a*::GFP knock-in feeding polyp (green) at day 0 post grafting. (**I**) Confocal images of a representative chimera at day 5 post grafting. White rectangles show close-up views of germinal zone (1) and sporosac (2), both composed of a mixture of GFP^+^ and GFP^−^ cells. Scale bar 100 μm.

Removing the head of feeding polyps results in regeneration of a new feeding head from the stump within 3 days^12,13^. Amputation of sexual heads, however, resulted in various outcomes, depending on the site of amputation. Amputating only the most distal (oral) part, leaving the germinal zone intact, resulted in the regeneration of a new sexual head (n=69/75). By contrast, if the entire oral part including the germinal zone were amputated, either a feeding head regenerated (n=11/75), or no head regenerated at all (n=64/75). In both cases, all sexual features degenerated (Figure S1). This suggests a pivotal role for the germinal zone in the maintenance of gonadal tissue identity.

Next, we studied the induction of sexual development. To do so, we conducted a series of experiments by heterotopic grafting of gonadal tissue onto feeding polyps (Figure 2B). We first grafted together two feeding polyps of the *Tfap2* reporter animal by joining their cut aboral ends, resulting in a bi-headed animal. This allowed one head to feed after removal of the other. We amputated one of the heads and replaced it with a new feeding head from a wild type donor. This resulted in the maintenance of the feeding identity of the chimeric animal with neither induction of somatic sexual structures (by morphology) nor germ cells (by lack of GFP) (Figure 2C).

We then replaced one of the feeding heads with a piece of the lower body of a wild type sexual polyp (i.e., a gonad) that contained immature sporosacs but no germinal zone. Normally, these sporosacs mature within days and spawn. However, the grafted sporosacs did not mature and gradually degenerated, nor did GFP^+^ cells appear, indicating an absence of somatic gonad development and of germ cell induction (Figure 2D). Next, we replaced one of the feeding heads with the oral tip of a wild type sexual polyp, again excluding the germinal zone. Within days, we observed in about half of them, the maintenance of the sexual head but no new sexual tissue was induced. In the other half, the sexual head was resorbed and a new feeding head grew instead. In both cases, no GFP+ germ cells were induced (Figure 2E).

Finally, when the germinal zone was included in the donor tissue, the sexual identity was maintained and new sporosacs developed. Moreover, *de novo* germ cells were induced in the recipient feeding polyp, attested by GFP expression (Figure 2F), showing that the germinal zone can not only maintain the sexual identity of the entire somatic gonad but also induce new germ cells. To identify possible induction of new somatic gonad tissue by the grafted germinal zone, we repeated the experiment using a different transgenic animal as recipient. This animal had GFP knocked-into its *Ef1a* gene^14^, resulting in ubiquitous GFP fluorescence (Figure 2G). We grafted a sexual head including the germinal zone from a wild type animal onto an *Ef1a* knock-in animal (Figure 2H). This resulted in the induction of both germ cells and somatic gonads in the recipient feeding polyp (Figure 2I).

Overall, the above experiments show that the germinal zone acts as a sexual organizer, being sufficient to induce and maintain sexual tissue that includes both somatic and germ cells in non-sexual tissue. The results also demonstrate that in the absence of a germinal zone, the feeding identity of polyps predominates, being the default fate of *Hydractinia* polyp development.

### *Gonadless* is a germ cell-specific member of the TGF-β family

To gain insight into the mechanisms of sexual induction, we reanalyzed a published differential gene expression dataset of different polyp types^11^. A gene, annotated as *Dvr1* in GenBank (accession No. HyS0012.146), is upregulated in the distal part of male and female sexual polyps. Phylogenetic analysis placed the gene in the TGF-β protein family that includes BMP, DVR, GDF, and NODAL proteins, but its orthologous relationship with vertebrate subgroup members could not be resolved (Figure S2). The TGF-β pathway is a major regulator of germ cell induction, maintenance, proliferation, and maturation in bilaterians^3^. The predicted protein has a secretory signal peptide in its N-terminus and a TGF-β domain in the C-terminal region. We hypothesized that this gene is involved in sexual development. For reasons outlined below, we named the gene *Gonadless* (*Gls*).

We performed Signal Amplification By Exchange Reaction (SABER) single molecule fluorescence mRNA in situ hybridization^15,16^ to study the expression pattern of *Gls*. These experiments showed that *Gls* mRNA is present at detectable levels exclusively in germ cells in the germinal zone that also express *Tfap2*, an early germ cell marker^11^ (Figure 3). Tfap2 is the master regulator of germ cell commitment, essential and sufficient to induce germ cell fate in pluripotent i-cells^11^. As shown previously, loss of *Tfap2* results in animals that have neither germ cells nor gonads^11^.

**Figure 3.**
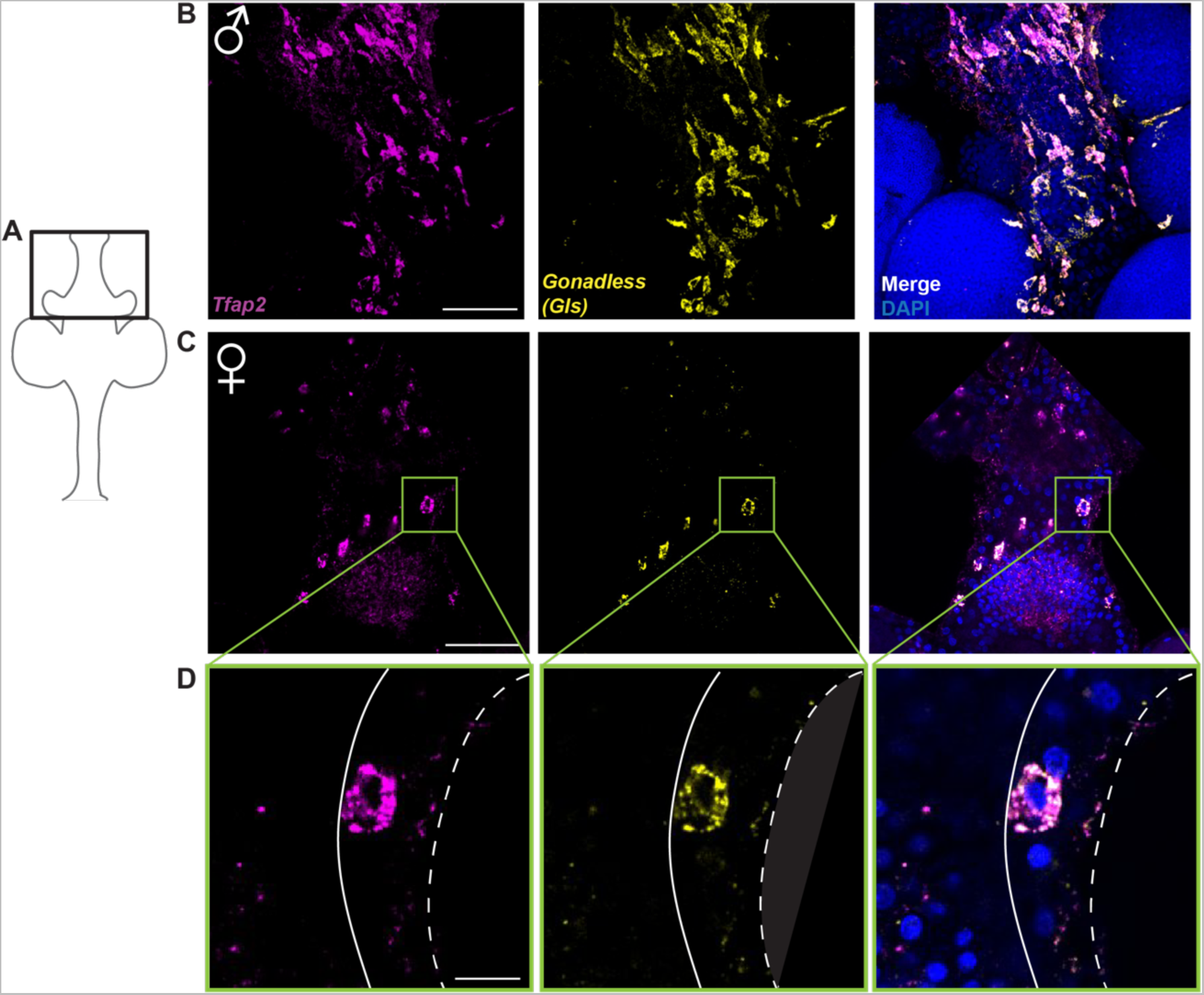
*Gls* expression pattern. (**A**) Cartoon of a sexual polyp. The black box corresponds to the position of the confocal images. (**B**) Maximum projection of mRNA in situ hybridization of *Tfap2* and *Gls* in a male sexual polyp. (**C**) Maximum projection of mRNA in situ hybridization of *Tfap2* and *Gls* in a female sexual polyp. Scale bar 50 μm. Green rectangle correspond to the close-up view shown in D. (**D**) Single confocal section showing expression of *Tfap2* and *Gls* in the cells boxed in C. White dashed line represents the outline of the epidermis and continuous white line represents the basement membrane (mesoglea) separating the gastrodermis from the epidermis. Scale bar 10 μm.

### Gls is required for gonad development

To address the function of Gls, we generated knockout animals using CRISPR/Cas9-mediated mutagenesis. Two single guide RNAs (sgRNA) were designed to flank the TGFβ domain of *Gls* (Figure 4A). We injected the sgRNAs together with recombinant Cas9 into zygotes and screened the G0 animals by PCR to identify individuals that had a genomic deletion in their cells. We grew the animals to sexual maturity and crossed them with wild type animals. Heterozygous G1 animals were selected by PCR and sequencing, grown to sexual maturity, and interbred to obtain homozygous *Gls*^−/−^ knockout animals. We obtained a total of eight knockout animals consistent with the predicted Mendelian distribution (Figure S3). These animals developed normally to planula larvae and metamorphosed to primary polyps, 24 hours post CsCl induction^17^. The animals developed to normal-appearing young colonies; however, none of them developed sexual polyps (i.e., gonads) even after 20 months (Figure 4B). Sexual polyps normally appear within 2 months in wild type *Hydractinia* colonies^18^. We concluded that Gls is essential to induce somatic gonad development (hence the name *Gonadless*).

**Figure 4.**
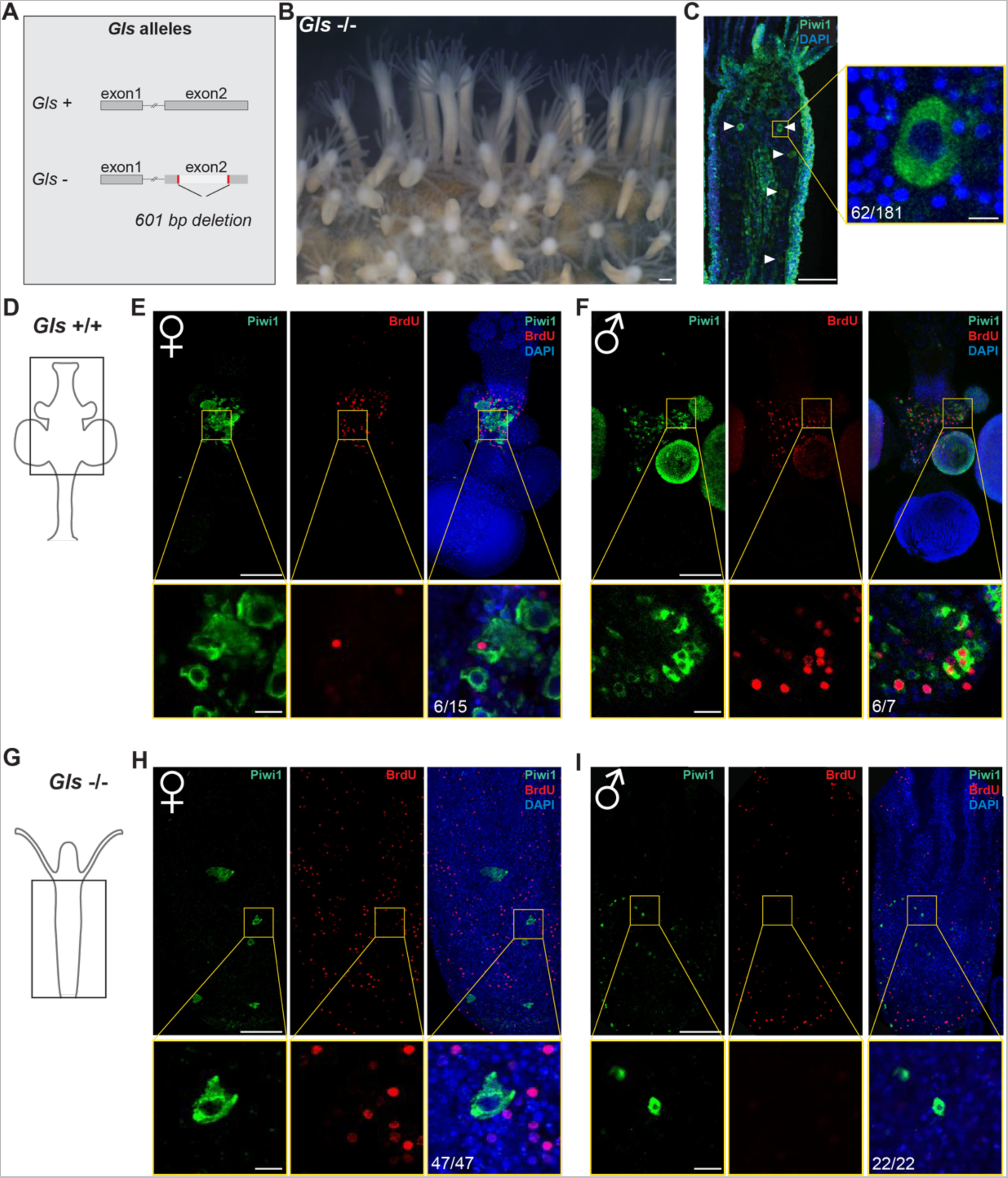
Morphological and cellular characterization of *Gls*^−/−^ animals. (**A**) Graphical representation of the genomic structure of wild type and mutant alleles of *Gls*. (B) A *Gls*^−/−^ mutant, having only feeding polyps. (**C**) Single confocal section of a *Gls*^−/−^ feeding polyp with Piwi1^+^ oocytes (white arrows) in its gastrodermis. Yellow box represents a close-up view. (**D**) Cartoon of a wild type sexual polyp. The black box corresponds to the position of the confocal images shown in E and F. (**E,F**) Maximum projection of immunostaining of Piwi1 (green) and BrdU (red) in the gastrodermis of a female (E) and a male (F). Yellow boxes represent a close-up view of a single confocal section shown below, respectively. Germ cells proliferate in wild type animals. (**G**) Cartoon of a Gls^−/−^ feeding polyp. The black box corresponds to the position of the confocal images shown in H and I. (**H,I**) Maximum projection of immunostaining for Piwi1 (green) and BrdU (red) in the gastrodermis of a female mutant (H) or of a male mutant polyp. Yellow boxes represent a close up of a single confocal section showing that germ cells do not proliferate in *Gls*^−/−^ mutants. Scale bar 100 μm. Scale bar in yellow boxes 10 μm.

### Gls is dispensable for primary germ cell induction

To gain insight into the cascade of sexual development, we looked for the presence of germ cells in *Gls*^−/−^ animals. We argued that if Gls acted directly upstream of *Tfap2*–the major regulator of germ cell fate–no germ cells should be present in *Gls* mutants. By contrast, the presence of germ cells in *Gls* mutants would be consistent with the gene being expressed downstream of Tfap2. We performed immunofluorescence (IF) using anti-Piwi1 antibodies on wild type and *Gls* knockout feeding polyps. Piwi1 is a marker of both pluripotent i-cells and germ cells. However, i-cells are only found in the interstitial spaces of the epidermis; they migrate soon after acquisition of germ cell fate to the gastrodermis where they mature to gametes. Therefore, epidermal Piwi1^+^ cells are mostly i-cells but Piwi1^+^ cells in the gastrodermis are exclusively germ cells.

In wild type feeding polyps, we did not find gastrodermal Piwi1^+^ cells (i.e., germ cells), except for one female polyp with a gastrodermal oocyte (Figure S4). However, in *Gls*^−/−^ feeding polyps, we found gastrodermal Piwi1^+^ cells in ∼34% in of the feeding polyps surveyed (Figure 4C and Figure S4). A 5–Bromo–2′–deoxyuridine (BrdU) incorporation followed by anti-BrdU and anti-Piwi1 immunofluorescence showed that these germ cells were not proliferative, in contrast to proliferative germ cells in wild type animals (Figure 4D-I). In one of the mutant clones, the gastrodermal Piwi1^+^ cells were oocytes (Figure 4C). The other mutants had small Piwi^+^ germ cells in their gastrodermis, leading us to believe that they were males (Figure S4). No mature eggs or sperm were found in *Gls*^−/−^ animals. Therefore, while Gls is required for the development of the somatic tissue of the gonad and, indirectly, for germ cell proliferation and maturation, it is not necessary for primary germ cell induction.

### Gls is essential for secondary germ cell induction

To rescue the sexual phenotype in *Gls* mutants, we grafted a sexual head including the germinal zone from a *Gls* wild type, *Ef1a* knock-in animal with ubiquitous GFP expression, onto a decapitated *Gls*^−/−^ knockout feeding polyp (Figure 5A). This experiment allowed the reintroduction of Gls signaling in the mutant and a direct observation of the outcome by fluorescence microscopy since the mutant cells were not fluorescent. It resulted in the induction of both somatic gonads and germ cells in the mutant (Figure 5B), showing that a cascade driven by wild type *Gls* expression is sufficient to induce full sexual development, including somatic gonadal tissue and secondary germ cells.

**Figure 5.**
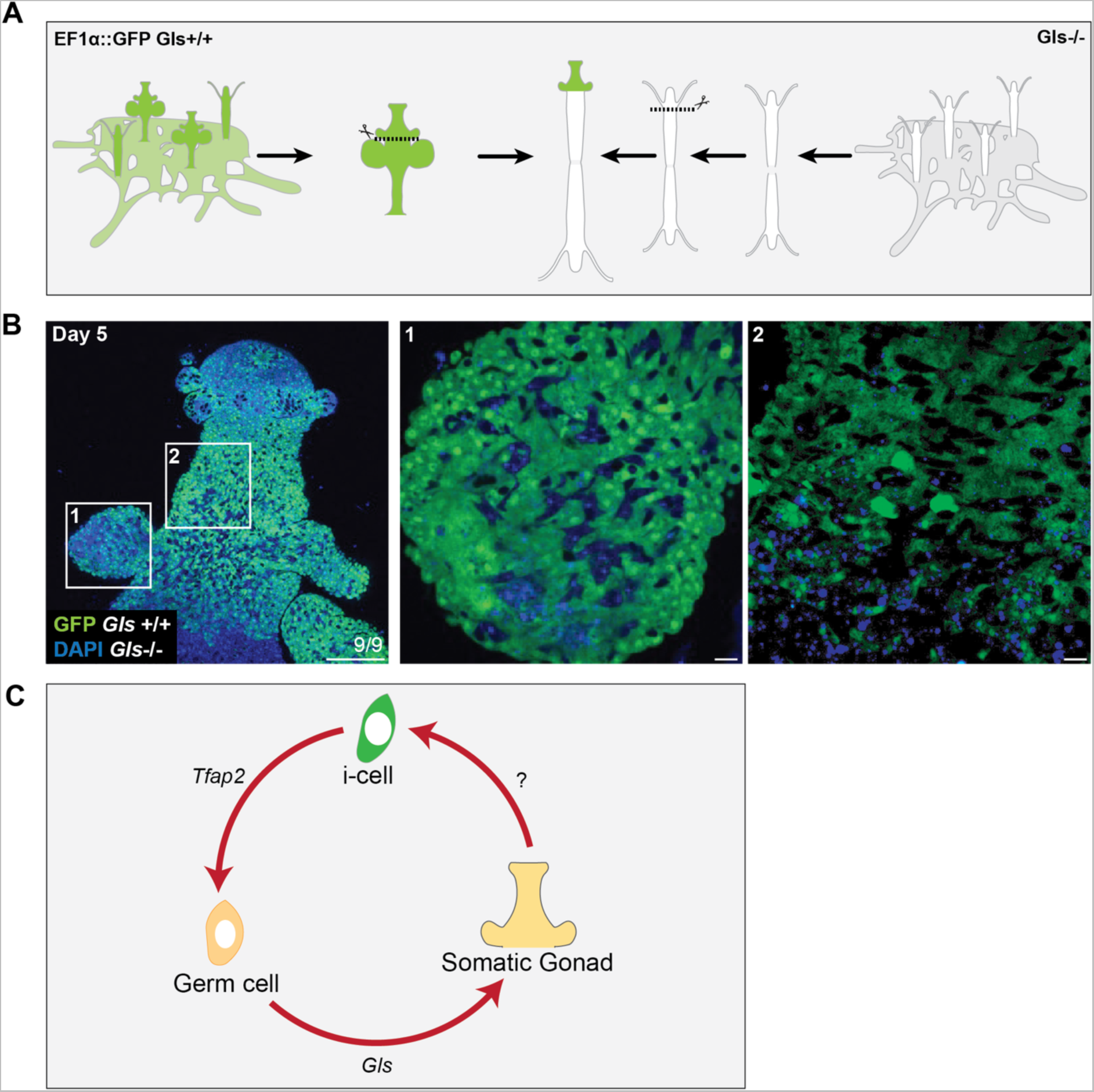
Heterotopic grafting of *Ef1a::*GFP fluorescent *Gls*^+/+^ sexual tissue onto sterile Gls^−/−^ animals. (**A**) Cartoon of the experimental grafting procedure. (**B**) Confocal images of a chimera, 5 days post grafting. Scale bar 100 μm. White squares represented closes-up images of a newly formed sporosac (1) and the germinal zone (2), showing that they are composed of a mixed of *Gls*^−/−^ (GFP^−^) and *Gls*+/+ (GFP^+^) cells. Scale bar 10 μm. (C) A model for *Hydractinia* sexual development. Germ cells induce somatic gonad morphogenesis via Gls signaling, and the newly formed tissue, in turn, promotes secondary germ cell induction (by activating *Tfap2* in pluripotent stem cells), proliferation, and maturation.

### Gls is insufficient for precocious sexual development

To probe the ability of ectopic TGF-β signaling to activate the sexual program, we forced the activation of downstream signaling events using the BMP agonist, sb4. We incubated intact and decapitated *Tfap2*::GFP reporter feeding polyps with sb4. No induction of sexual somatic tissue or germ cells was observed in the treated animals. However, they tended to grow stolon and settle on the bottom of the Petri dish or adhere to each other (Figure S5). Finally, we generated transgenic animals that expressed *Gls* ectopically in several contexts. Given the absence of conditional expression protocols for *Hydractinia*, we used the promoters of several genes to drive *Gls* in embryos. First, we used the genomic control elements of the *Rfamide precursor* gene to drive *Gls* expression in embryos. *Rfamide precursor* is expressed in a subset of neurons in the heads of feeding and sexual polyps^13^. The transgenic construct also included GFP, separated by a T2A peptide sequence to mark transgenic cells in the mosaic animal. Most transgenic animals died during embryogenesis or before completing metamorphosis and the remaining ones developed with severe patterning defects (Figure S6). We repeated the experiment using different genetic elements to drive *Gls* in *Tfap2* reporter animals: *Wnt3* is expressed in the oral tips of polyps (Figure S6); *Ncol1*, a nematogenesis marker, is expressed by differentiating nematocytes at the lower part of the body column (Figure S6). Finally, we used the *B-tubulin* genomic control elements to drive *Gls* ubiquitously (Figure S6). All these constructs generated various, mild to severe defects in development but no germ cells were induced as evidenced by lack of GFP reporter fluorescence (Figure S6). Only in rare cases did we observed structures that resembled sexual polyps in metamorphosed animals (Figure S6). Therefore, Gls is insufficient to induce sexual development precociously.

## Discussion

*Gls*, a member of the TGF-β family, is expressed by germ cells and required for patterning and maintenance of the somatic gonadal tissue. We also demonstrate that the germinal zone within the gonad, in turn, facilitates the induction of new germ cells through activation of *Tfap2* in pluripotent i-cells.

We propose a model for sexual development in *Hydractinia*. In the model, a small number of germ cells are induced, by an unknown factor, in some polyp buds in sexually mature colonies. We call these primary germ cells. These germ cells secrete Gls protein that induces the development of the germinal zone that acts as a sexual organizer. The germinal zone secretes a factor that induces *Tfap2* expression in resident pluripotent i-cells, committing them to germ cell fate that we call secondary germ cells. The gonadal somatic tissue also provides the microenvironment that promotes proliferation of germ cells and eventually their maturation to gametes. Secondary germ cells, in turn, secrete Gls to maintain the germinal zone and by proxy, the entire gonad. Therefore, a positive feedback loop induces and maintains sexual reproduction (Figure 5C).

Several questions remain unanswered. First, what is the nature of the signal that induces primary germ cells? Second, what is the signal that is emitted from the germinal zone in the somatic gonad to induce *Tfap2* expression in pluripotent i-cells and thereby convert them to secondary germ cells? The possibility that Gls itself acts on i-cells is unlikely since precocious expression of Gls was insufficient to induce germ cells. Moreover, if this were the case, germ cells would induce all i-cells in their vicinity to become germ cells too and germ cell identity would spread uncontrollably. Therefore, Gls induces somatic gonad patterning, but the signal emitted by the gonad to induce secondary germ cells must be distinct from Gls to restrict secondary germ cell commitment to the germinal zone. Finally, what molecular signals act downstream of Gls and induce proliferation of germ cells and their maturation to gametes?

It is interesting to speculate about the evolution of the germ cell-gonad interactions. Germ cells in all studied animals are specified before the development of gonads. Therefore, it is possible that induction of gonad morphogenesis by a TGF-β signal of germ cell origin is a primitive trait in animals. By contrast, since most bilaterians segregate a germline only once in a lifetime, induction of secondary germ cells in the gonad does not exist in these animals. It has either been lost in bilaterians or constitutes a cnidarian innovation.

## Materials and methods

### Animals

*Hydractinia symbiolongicarpus* colonies were kept in artificial seawater (ASW) as described^9^. Fertilized eggs were collected 1.5 hours after exposing male and female colonies to light.

### Grafting

*H. symbiolongicarpus* colonies were starved for a day and anesthetized in 4% MgCl_2_ (in 50% distilled water/50% filtered seawater). Polyps were dissected from the colony. Two feeding polyps were grafted at their wound side by skewering them onto a minutien pin (Fine Science Tools, CAT#26002-10). They were then pressed together by placing a block of 1% agarose gel at each end and left for 2 hours for the tissues to fuse. Grafts were removed from the minutien pin by gently pushing them toward the end of the pin with tweezers. Grafts were transferred to a glass Petri dish with freshly 0.2 μm filtered seawater and incubated overnight. The next day, one head of the grafts was removed. Donor tissue feeding heads or sexual polyp parts (oral tip, germinal zone, mature sporosac) were isolated using a scalpel. Donor and recipient tissues were skewered onto an minutien pin. They were then pressed together by placing a piece of 1% agarose gel at each end and left for 4 hours for the wound to heal. Chimeras were removed from the minutien pin and transferred to a glass Petri dish with freshly 0.2 μm filtered seawater and incubated with agitation for 8 days at 18°C. Chimeras were fed with *Artemia franciscana nauplii* and seawater was replaced daily.

### BrdU incubation

Sexual polyps were incubated for one hour in 150 mM BrdU (Sigma; Cat. #B5002) in ASW and fixed in 4% paraformaldedhyde/ASW.

### Injection of embryos

One-cell stage embryos of wild type or *Tfap2* reporter animal were injected using a Narishige IM 300 microinjection system as described^11^.

### Ectopic expression of *Gls*

*Gls* coding sequence in-frame with T2A peptide was synthetized by IDT. Insert was ligated into the 5’Wnt3::GFP::3’Wnt3^11^ and 5’RFamide::GFP::3’*β*tubulin^13^ using restriction enzymes Not1 and Sac1. Insert was ligated into the 5’*β*tubulin::mScarlet::3’*β*tubulin (Dubuc and al., 2020) and 5’Ncol1::mScarlet::3’Ncol1^19^ plasmids using Gibson assembly according to the manufacturer recommendations (NEB, CAT#E2611). Sequences of *Gibson* primers used were:

*β*tubulin-Fwd CCAGGTCCAATGGTATCTAAAGGTGAAGC;
*β*tubutulin-Rev ACGATGACATTTTTTCAGATCCACCTCC;
*β*tubutulin-Gls-T2A-Fwd ATCTGAAAAAATGTCATCGTTGTTAATATTTTTG;
*β*tubutulin-Gls-T2A-Rev TAGATACCATTGGACCTGGGTTTTCTTC;
*Ncol1-* Fwd CCCAGGTCCAATGGTATCTAAAGGTGAAGC;
Ncol1-Rev ACGATGACATTACTGTAGGATTGTTATAATCG;
Ncol1-Gls-T2A-Fwd TCCTACAGTAATGTCATCGTTGTTAATATTTTTG;
Ncol1-Gls-T2A-Rev TAGATACCATTGGACCTGGGTTTTCTTC.

The full sequence of the plasmids injected can be found in Supplemental file.

Plasmids listed were injected at 500-1800 ng/µl and were supplemented with 200 mM of KCl.

The phenotype of the animal was assessed at 2 days after metamorphosis.

### CRISPR/Cas9 knock-out

Short guide RNAs (sgRNAs) were designed using Geneious (2017.9.1.8). The three selected sgRNAs for *Gls* did not match other genomic sequence. Synthetic sgRNAs were provided by Synthego Inc and diluted according to the manufacturer’s recommendations. All sgRNAs were incubated together (500 ng/µl) with recombinant Cas9 (1 μg/μl; IDT, Cat. #1074181) for fifteen minutes prior to injection as described in^16^.

### Genotyping

Forty-five CRISPR-Cas9 injected animals were metamorphosed. Three animals displayed defects in sexual development. Animals were genotyped for *Gls* mutations. Primers spanning the entire second exon of the gene (see FigS3 and Supplemental file) were used in a PCR reaction to identify large deletions. PCR products were sequenced, allowing us to identify a 601bp deletion in Mutant 28. Sequence of the *Gls* mutant 28 used for the generation of G1 heterozygotes can be find in Supplemental file. G1 animals were metamorphosed and grown to sexual maturity. G1 males and females that carried the 601bp deletion were then crossed to obtained G2 offspring. G2 animals were genotyped, and eight homozygote mutant animals were identified.

DNA was extracted as described in^16^. A two-step PCR approach was used to amplify genomic fragments. PCR conditions using Phusion High-Fidelity DNA Polymerase (Thermo Fisher Scientific, Cat. #F530S,) were as follows: initial denaturation (98°C, 3:00), denaturation (98°C, 0:30), annealing/extension (64°C, 2:30), repeat steps 2-3 (30x), and a final extension (64°C, 15:00). PCR reactions were run on a 1% agarose gel and positive bands were excised and purified using Monarch Gel extraction clean kit (NEB, Cat. #T1020L). A-tailing of the fragments was carried out by incubating the DNA with MyTaq polymerase (NEB, Cat. #M0273) for 15 minutes. Inserts were then ligated into the pGEM-T Easy Vector System (Promega, Cat. #A1360). DH5-alpha *E. coli* bacteria were transformed and plated on ampicillin LB-agar plates and grown overnight. Individual clones were picked and grown in overnight cultures of LB-broth (with ampicillin, 100µg/ml). Plasmid DNA was extracted using the Monarch Miniprep Kit (NEB, Cat. #T1010L) and sent to sequencing to identify mutations.

### In situ hybridization

Probes were synthesized according to Kishi et al (2019)^15^ using hairpins 27 and 30. Tissue was fixed and dehydrated as previously described in^11^. In situ hybridization experiments were performed as described previously^16^, with changes listed below. After rehydration, samples were incubated for 10 min in proteinase K (2mg/ml) in PBS Tween 0.1% and fixed in 4% paraformaldedhyde/PBS.

### Immunofluorescence

Immunofluorescence staining was performed as previously described^20^. Anti-Piwi1 antibodies were used at the concentration of 1:2000^11^; anti-GFP 1:300 (Synaptic Systems, CAT#132005); 1:100 anti-BrdU (Abcam, CAT#ab6703).

### Phylogeny

Putative *Hydractinia* genes were identified with tBLASTn and BLASTp searches using human sequences as queries. Reciprocal BLAST was performed, including newly identified *Hydractinia* sequences. Newly identified genes from *Hydractinia* were next used as queries as well. After identification of genes in *Hydractinia*, proteins were analysed with ScanProsite (ExPaSy)^21^ to verify the presence of TGF specific domains.

Data set was built by collecting all sequences from a previous phylogenetic analysis of TGF family^22^ to which we added *Hydractinia* sequences. In addition, we completed the dataset by incorporating additional sequences from selected species, especially cnidarians *Hydra vulgaris*, lophotrochozoans *Biomphalaria glabrata*, cephalochordate *Branchiostoma floridae*, and teleost *Danio rerio* in order to build an alignment more representative of metazoan diversity. From these species we added Nodal and TGF beta genes which were absent/underrepresented in the original alignment despite them belonging to the TGF family^22^. Multiple sequence alignment of amino acids was generated using MAFFT version 7^23^ with default parameters. Gblocks version 0.91b^24^ was used to remove vacancies and blur sites. Final alignment is comprised of 376 amino acids. Multiple sequence alignment can be found in Supplemental file 1.

Phylogenetic analyses were run from the amino-acid alignment using maximum-likelihood method with PhyML 3.1 software^25^. Analysis was carried out under the WAG evolution model, maximum likelihood was assessed using MEGA11^26^. Node robustness was evaluated by conducting 500 bootstrap replicate sampling.

Bayesian analyses were performed using MrBayes (v3.2.6)^27^ under mixed model. Similarly, the WAG evolution model was determined to by the most likely under the Bayesian method. The phylogenetic analysis was carried out for 300,000 generations with 10 randomly started simultaneous Markov chains (1 cold chain, 9 heated chains) and sampled every 100 generations. One fourth of the topologies were discarded as burn-in values, while the remaining ones were used to calculate posterior probability. Node robustness corresponds to posterior probabilities.

For both phylogenetic analyses the outgroup is the human GDNF sequence (NCBI P39905.1) already successfully used to conduct TGF family phylogeny^22^.

### Imaging

Images of whole colonies and chimeras were taken using a Leica M165FC microscope with a Leica DFC7000T camera. Confocal images were acquired using an Olympus Fluoview 3000 laser scanning confocal microscope. Images were imported into ImageJ/Fiji. Their brightness and contrast were adjusted as a whole. To facilitate visualization, a Gaussian blur correction was applied to the images of the SABER *in situ*. We aimed to display all polyp images in the same orientation, therefore some images were rotated, and a black rectangle was inserted behind for aesthetic reasons.

## Acknowledgements

We thank members of the Frank lab for inspiration and discussion. All microscopy work was conducted at the Microscopy and Imaging Core Facility of the University of Galway. This work was funded by a Wellcome Trust Investigator Award to UF. GK was an Irish Research Council postdoctoral fellow; PKW is an Irish Research Council postgraduate fellow.

## Author contribution

CC, TQD, and UF conceptualized the study. CC performed experiments; CD and CC performed immunofluorescence; PKW and CC performed mutant screening; GK and HRH performed Gls phylogeny; CC and UF wrote the paper.

---

**Figure S1.**
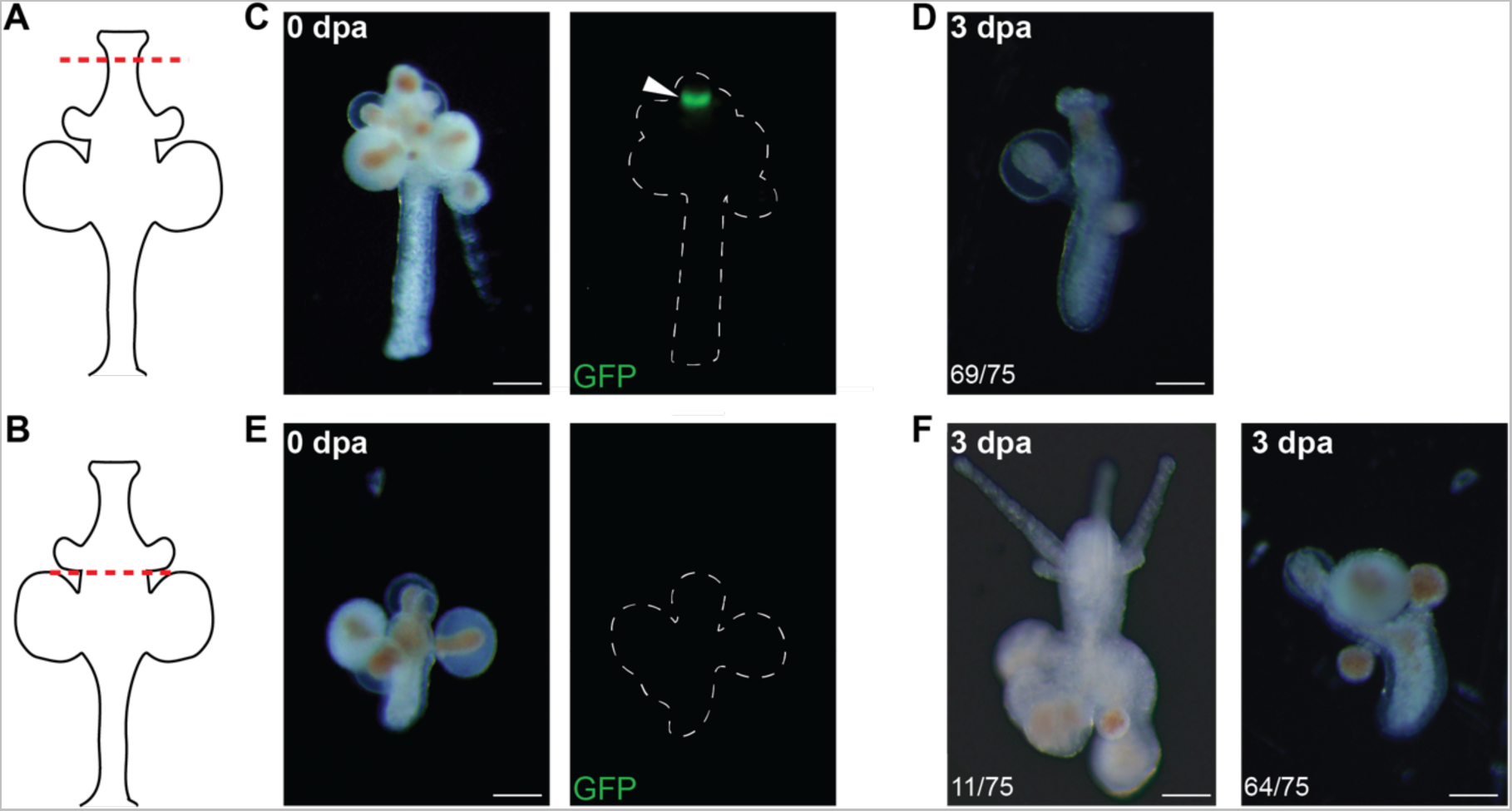
Sexual head regeneration. (**A, B**) Cartoon of a sexual polyps. Red dashed lines indicate amputation planes for the experiments shown in C and D, and E and F, respectively. Pictures of regenerating *Tfap2*::GFP polyp 0 days post amputation. White dashed lines show the outline of the polyp in the GFP channels. Scale bars 100 μm.

**Figure S2.** Gls phylogeny. Topology of TGF multigene family relationships obtained by phylogenetic analysis using Bayesian inference (**A**) and maximum likelihood (**B**) methods on amino acid sequences. The phylogenetic trees are rooted by the human gene GDNF (Glial Derived Neurotrophic Factor). The two analyses gave rise to similar topologies with two major groups which are the TGF-β clade and the Decapentaplegic-Vg1 related (DVR) family clade. In both, the activin/inhibitin sequences present a paraphyletic distribution close to the root. Even if some variation can be observed, the deep nodes remain conserved. The DVR related family is subdivided between BMPs, Nodal, and a group composed of both GDF and BMP. The *Hydractinia* genome encodes nine TGF-β multigene family members, including two actinin/inhibitin, three BMP, three GDF/BMP related genes, and Gonadless. *Hydractinia* sequences are labelled by white arrows, expect Gonadless pointed with black arrow. (Bf: *Branchiostoma floridae*; Bg: *Biomphalaria glabrata*; Dr: *Danio rerio* Dm: *Drosophila melanogaster*; Cg: *Crassostrea gigas*; Ed: *Exaiptasia diaphana*; Ga: *Gigaantopelta aegis*; Gg: *Gallus gallus*; Hr: *Halocynthia roretzi*; Hs: *Homo sapiens*; Hv: *Hydra vulgaris*; Hsy: *Hydractinia symbiolongicarpus*; Mm: *Mus musculus*; Sp: *Strongylocentrotus purpuratus*; Xl: *Xenopus laevis*). Nodes robustness are posterior probabilities and bootstraps (500 replicates) for Bayesian and maximum likelihood analyses, respectively.

**Figure S3.**
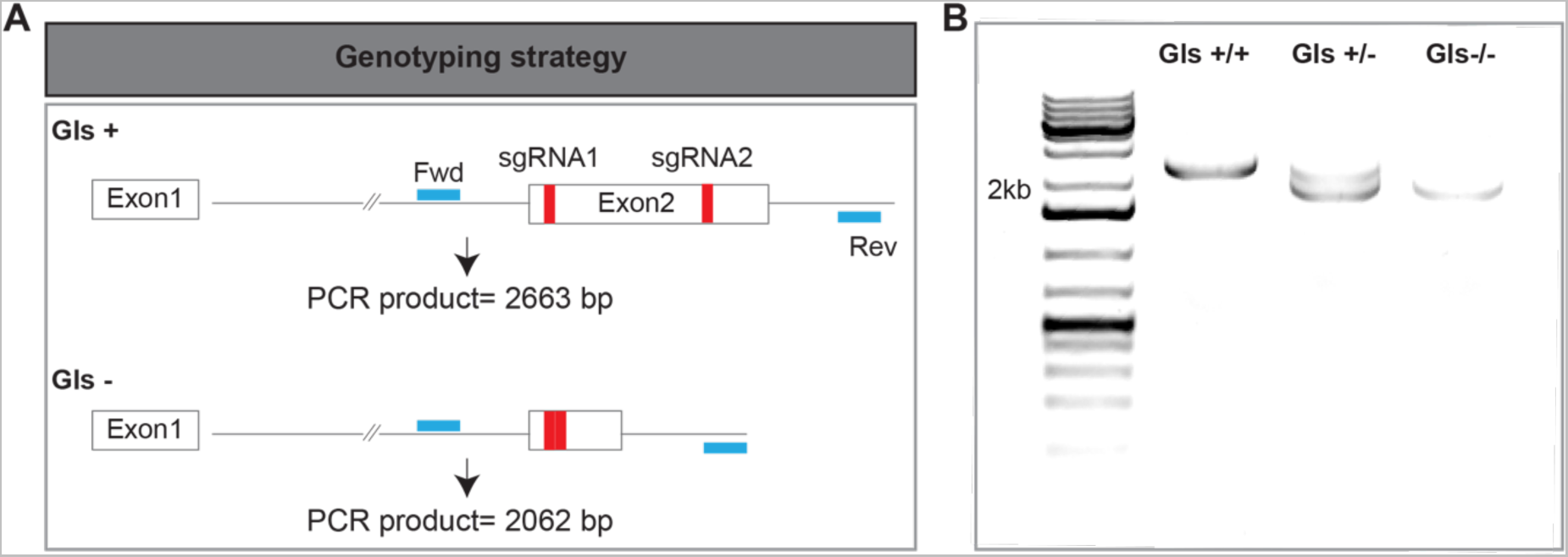
Screening for genotyping of *Gls* mutants. (**A**) Graphical representation of the wild type and mutated alleles and the screening strategy. Red bars show the position of the Cas9 cut sites. (**B**) Results of the PCR genotyping showing wild type, heterozygous mutant, and knockout homozygous mutant.

**Figure S4.**
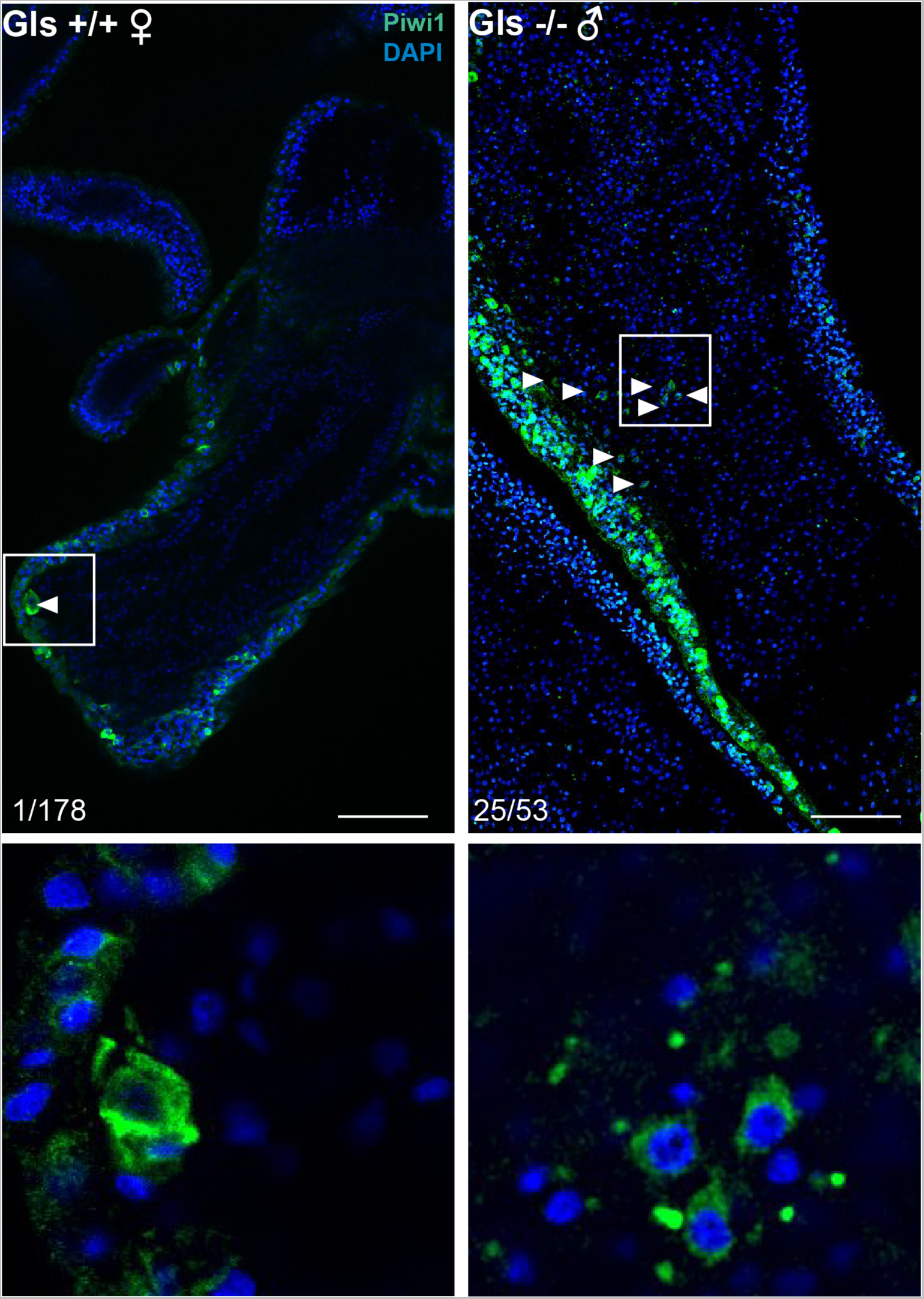
Piwi1^+^ cells in wild type and Gls^−/−^. Confocal images of a female Gls^+/+^ and a male Gls^−/−^ showing Piwi1^+^ germ cells (white arrows) in their gastrodermis. White boxes represent a 50 μm close-up of these cells. Scale bar 50 μm.

**Figure S5.**
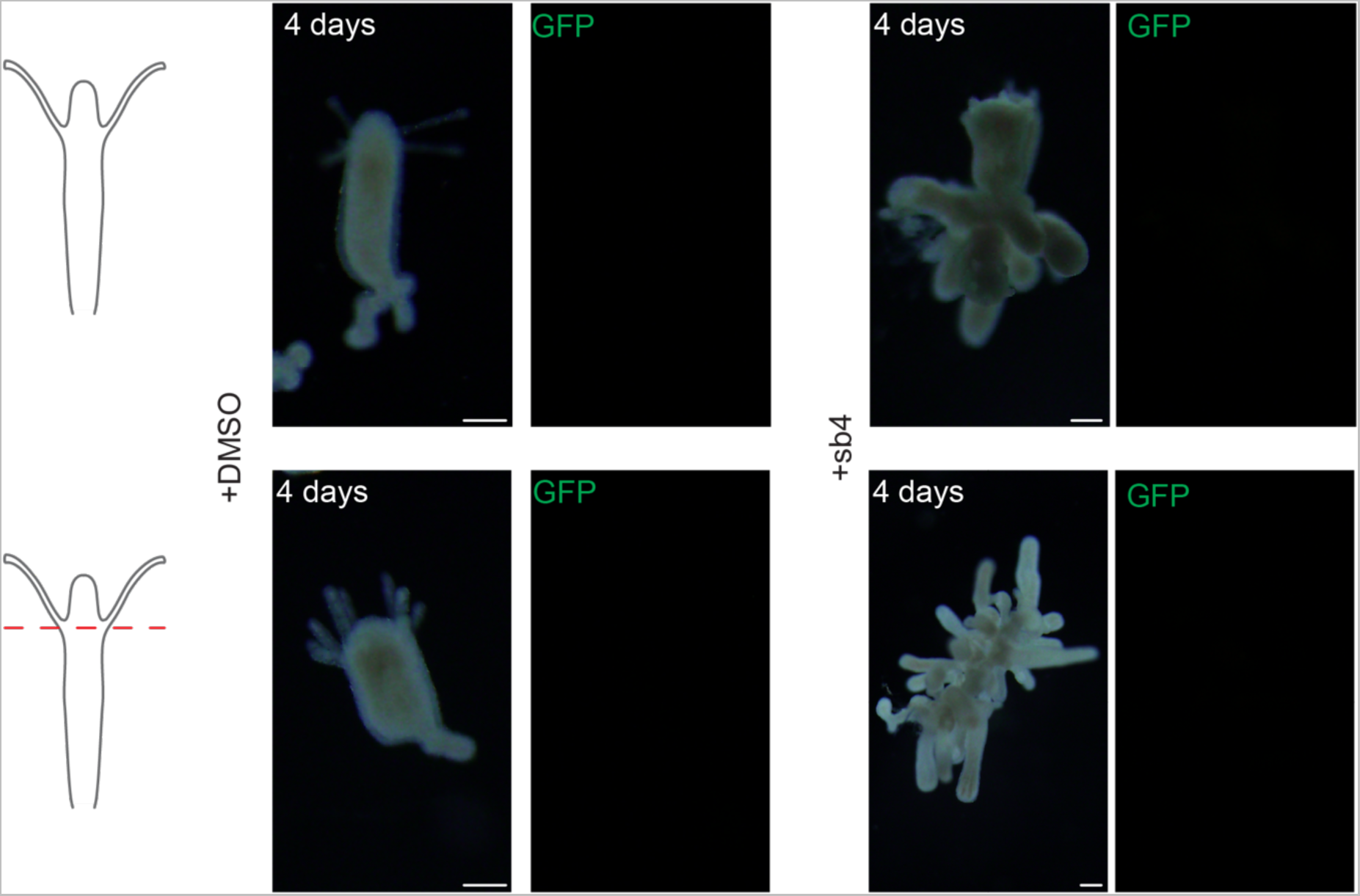
Head regeneration with BMP agonist. Intact and decapitated *Tfap2*::GFP feeding polyps incubated with DMSO (control) or sb4 (BMP agonist) at 1 μM showing that no somatic sexual structures or germ cells (GFP) were induced 4 days upon treatment. N=25 in each condition. Red dashed line shows the site of amputation. Scale bar 100 μm.

**Figure S6.**
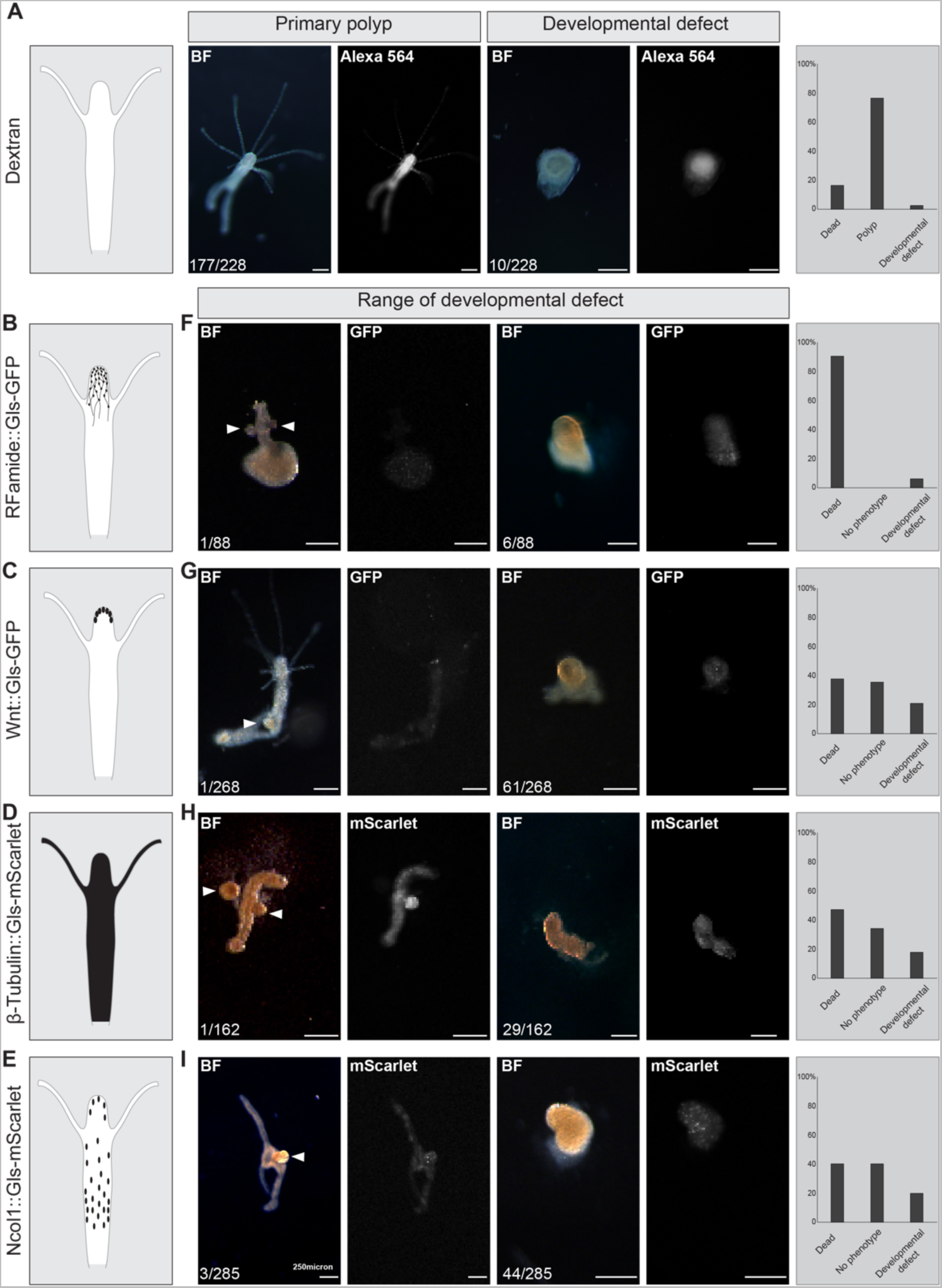
Ectopic expression of Gls in embryos. (**A**) Control embryos injected with dextran 565 develop into a primary feeding polyp. (**B**) *Rfamid precursor* promoter-driven Gls-GFP. (**C**) *Wnt3* promoter-driven Gls-GFP (**D**) *B-tubulin* promoter-driven Gls-mScarlet injected in *Tfap2*::GFP reporter animals (**E**) *Ncol1* promoter-driven Gls-mScarlet injected in *Tfap2*::GFP reporter animals. No germ cells (GFP^+^ cells) were detected (**F-I**) Transgenes expression cause mild to severe developmental defects. White arrows indicated structures that resembled sexual tissue in metamorphosed animals. Scale bar 250 μm. BF: bright field.

